# High throughput single cell sequencing of both *T-cell-receptor-beta* alleles

**DOI:** 10.1101/320614

**Authors:** Tomonori Hosoya, Hongyang Li, Chia-Jui Ku, Qingqing Wu, Yuanfang Guan, James Douglas Engel

## Abstract

Allelic exclusion is a vital mechanism for the generation of monospecificity to foreign antigens in B- and T-lymphocytes. Here we developed a high-throughput barcoded method to simultaneously analyze the VDJ recombination status of both mouse *T cell receptor beta* alleles in hundreds of single cells using Next Generation Sequencing.

## INTRODUCTION

Vertebrates have evolved both innate and adaptive immune systems to protect individuals against infection, cancer and invasion by parasites. B and T lymphocytes comprise the central components of adaptive immunity and every individual lymphocyte harbors specific reactivity to a single antigen that is conferred by individually unique antigen receptors expressed on the cell surface. The diversity of antigen receptors is generated by DNA recombination, called VDJ rearrangement in jawed vertebrates(1). To maintain the required monospecificity of mature lymphocytes, only one of the two autosomal alleles is allowed to express a functionally rearranged beta- and alpha- chain T cell receptors (TCR*β* and TCR*α*) in T cells, or immunoglobulin heavy and light chain receptor (IgH and IgL) in B cells: both are controlled by a historically opaque mechanism referred to as “allelic exclusion”. Allelic exclusion occurs at the genetic level for IgH, IgL and TCR*β*(2), while at the level of protein localization on the cell surface for TCR*α*(3). Loss of allelic exclusion results in dual-TCR expression, which can lead to autoimmunity(4, 5).

A high-throughput method to study the diversity of lymphocytes at the single-cell level has been reported recently(6), yet how one might analyze their mono-specificity remains unclear. Since the abundance of transcripts from a non-functional allele is lower than that of the functional allele(7, 8), traditional RNA-based methods cannot detect the mechanisms underlying allelic exclusion. One major challenge in analyzing the genes themselves is that there is only one copy of DNA representing each allele, unlike the existence of multiple transcribed RNA species. Therefore, extremely accurate DNA sequencing is required to avoid erroneously mis-assigning nucleotide-level mutations, which would alter the interpretation of TCR locus activity. Additionally, no reference genome is available for merging multiple sequencing reads, due to the many millions of possible sequences that can be generated from VDJ rearrangement(9). Thus, *de novo* sequence assembly or a long-read approach is required to retrieve the original genomic sequence of both alleles in single cells. To significantly improve the traditional Sanger sequencing method employed by us and others(10-12), we developed a high-throughput method that enables analysis of *Trb* allelic exclusion status by sequencing both alleles of the genome in single cells, enabling us to determine whether each allele in those cells underwent either no, unproductive or productive rearrangement.

## MATERIALS AND METHODS

### Single cell isolation

Staged thymocytes were first isolated from mice (C57BL/6J, Jackson Laboratories) between 5 and 8 weeks old using a cell sorter (BD FACSAria III). Lin^-^CD4^-^CD8a^-^Thy1.2^+^cKit^-^CD25^+^CD28^-^ (DN3a stage), Lin^-^CD3^-^TCR*β*^-^ CD8a^-^ Thy1.2^+^cKit^-^CD25^-^(DN4 stage), CD4^+^CD8^+^(DP stage) and TCR*β*^+^CD4^+^CD8^+^(late DP stage) thymocytes (10,000 to 100,000 cells at each stage) were isolated as described previously(10). Next, single cells were directly sorted into 20 μl of lysis buffer [containing 1x Q5® Reaction Buffer (NEB), 4 μg Proteinase K and 0.1% Triton X100] in one well of a 96 well PCR plate using a Synergy cell sorter (Sony iCyt SY3200). The cell sorter setting was carefully aligned so that sorted cells were precisely deposited into the center of each well. Sorted single cells in the lysis solution were kept on ice, and then digested at 55°C for 60 minutes followed by 95°C for 15 minutes (to inactivate Proteinase K) using a PCR thermal cycler within 6 hours after sorting.

### Multiplex nested PCR

For the first round of PCR, primers were selected to amplify all potential VDJ rearrangement at the *Trb* locus: 31 V region primers covering all 35 *Trbv* genes(13), 2 D region primers, 2 J region primers and 2 control primers used to detect sequences 3’ of the *Actb* gene (Figs. 1a and 1b). The full list of primers used are deposited at (https://umich.box.com/s/3f5q64u2i68dn2i7hlneucxph9oetpvb). Since this method was designed to recover not only rearranged but also germ line configured genomic DNA, the two J primers were designed to be 3’ of J1-7 and J2-7. The following PCR condition amplifies the entire J1 (J*β*1 primer coupled with a V or D region primer) as well as J2 genomic DNA region (J*β*2 primer coupled with a V or D segment primer).

**Fig. 1.**
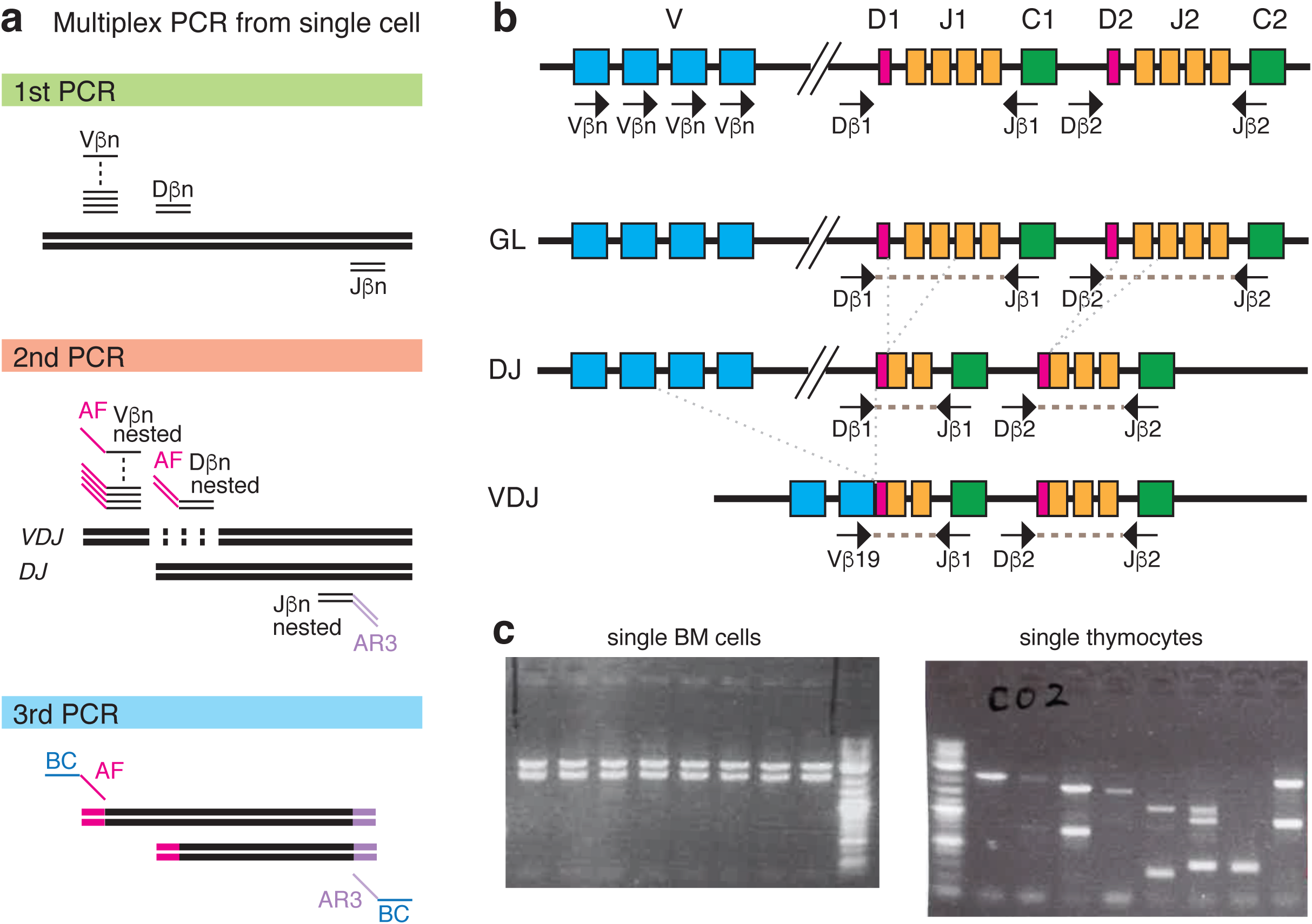
High-throughput *Trb* genomic DNA sequencing from single-cells. (**a**) Strategy for multiplex PCR amplification of genomic DNA sequences at the *Trb* gene VDJ loci from single cells. Final PCR products were mixed and then sequenced by PacBio RS II sequencer. V*β*n: V region primers. D*β*n: D region primers. J*β*n: J region primers. AF: adapter forward. AR: adapter reverse. BC: barcode. Details are described in Methods. (**b**) Schematic presentation of VDJ rearrangement at the *Trb* locus with simplified illustration of multiplex primer location. Genomic DNA recombination events are initiated by recombining D*β* (diversity) and J*β* (joining) segments on both chromosomes. Subsequently, on only one chromosome, one of the V*β* (variable) segments is joined to the previously rearranged DJ recombinant. Not only the selection of each VDJ segment, but also multiple lengths of spacer sequence between V, D and J, generates an incalculable number of VDJ sequence possibilities. mRNA splicing joins C*β* (constant) segments to the rearranged VDJ DNA recombinants to generate a final TCR*β* protein. VDJ rearrangement that generates a stop codon or elongated transcript results in a predictedly unproductive TCR*β*. (**c**) Successful amplification of rearranged VDJ, DJ and germline D-to-J regions were confirmed by agarose gel electrophoresis with NEB 2-log DNA ladder in the last (left panel) and first (right panel) lanes.

Since VDJ will be spliced to a C region at the RNA level, the selected J*β* primer sequences remain in genomic DNA after recombination. D region primers were designed 5’ of D1 and D2. The D segment primers amplify only a D-to-J rearranged genome, but not a V-to-DJ rearranged genome. After V-to-D1J rearrangement, the D1 primer sequence is removed from the genome. Similarly after V-to-D2J2 rearrangement, the D2 primer sequence is removed from the genome. The first round of PCR was performed in a 60 μl final reaction volume containing of 50 nM of each primer, 1x Q5® Reaction Buffer, 200 μM each dNTPs, 0.4 unit Q5® High-Fidelity DNA Polymerase (NEB) and 20 μl of the lysed single cell solution. The PCR condition was 30 sec at 98 °C followed by 30 cycles of 5 sec at 98 °C, 10 sec at 66-58 °C and 2 min at 72 °C, and then final extension for 2 min at 72 °C. During the first 5 cycles, the annealing temperature was reduced 2 °C per cycle from 66 °C to 58 °C, and then performed at 56°C for the last 25 cycles.

For the second round of PCR, nested primers were selected: 32 nested V region primers containing forward adapter sequences (AF-V*β*n), 2 nested D region forward adapter primers (AF-D*β*n) and 2 nested J region primers in a reverse adapter orientation (AR3-J*β*n). The second round of PCR was performed in a 20 μl final reaction volume containing 50 nM of each primer (Fig. 1a), 1x Q5® buffer, 200 μM of each dNTP, 0.4 units Q5® High-Fidelity DNA Polymerase (NEB) and 1 μl of the first round PCR product. The 2nd PCR condition was same as for the first round PCR but for only 25 cycles.

Finally, in the third round of PCR, unique barcoded-AF and barcoded-AR3 combinations of primers were selected for each single cell (Fig. 1a). Barcodes were adopted from a published report(6). The 3rd PCR condition was same as for the second round PCR (5 + 20 cycles). Hot start PCR was performed for all PCR reactions using either a BioRad T100™ or Applied Biosystems 2720 Thermal Cycler. All primers were purchased from Integrated DNA Technologies, Inc. The 37 (for 1^st^ round) or 36 (for 2^nd^ round) PCR primers at 200 μM final concentrations were mixed, aliquoted and frozen for subsequent use. Each PCR mix was prepared immediately before initiating PCR reactions. (The amplification efficiency was reduced if the PCR mix was prepared in advance and repeatedly frozen and thawed. Amplification efficiency was also reduced when 500 nM of each primer was used in the 2nd round PCR reaction). Successful DNA amplification was confirmed by running 2 μl of 3rd round PCR product on agarose gels with 0.4 μg of 2-Log DNA Ladder (NEB) (Fig. 1c): 8 samples were selected from each 96 well PCR plate. Of note, the PCR conditions employed here allow amplification of unrearranged (germline configuration) genomic DNA (Fig. 1c left panel).

### Recovery frequency

To analyze the frequency of recovered wells, a primer pair amplifying the 3’ region of the *Actb* gene was added to the 1st round PCR. 1 μl of the 1st round PCR product was used in a PCR reaction with nested *Actb* primers: 5’-CATAGGCTTCACACCTTCCT-3’, 5’- CTTTGCCTCCATCTGCATAAC-3’ and FAM-labeled probe TGCTAGTCTGAAGCTGCCCTTTCC (ZEN / Iowa Black FQ) purchased from Integrated DNA Technologies, Inc. Luna Universal Probe qPCR Master Mix (NEB, M3004) was used in a 20 μl reaction. The PCR condition was 30 sec at 95 °C followed by 45 cycles of 1 sec at 95 °C and 20 sec at 60 °C using StepOnePlus™ Real-Time PCR System (Thermo Fisher Scientific). A 90-95% recovery efficiency was routinely obtained from each plate containing 96 single cell sorted wells.

### PacBio high-throughput sequencing

Since the length of the PCR products described above ranged from 200 to 2700 bp (Fig. 1b), a sequencer with the ability to read long DNA fragments is preferred. To this end, Pacific Biosciences single molecule real-time (SMRT) sequencing technology(14) was employed. A PacBio smartbell adapter was ligated to the barcoded PCR product mixture and then sequenced on PacBio RSII sequencer at the University of Michigan Sequencing Facility. Thus each circular consensus sequencing (CCS) read generated from PacBio sequencing corresponds to a single molecule of initial PCR product (Fig. 2a).

**Fig. 2.**
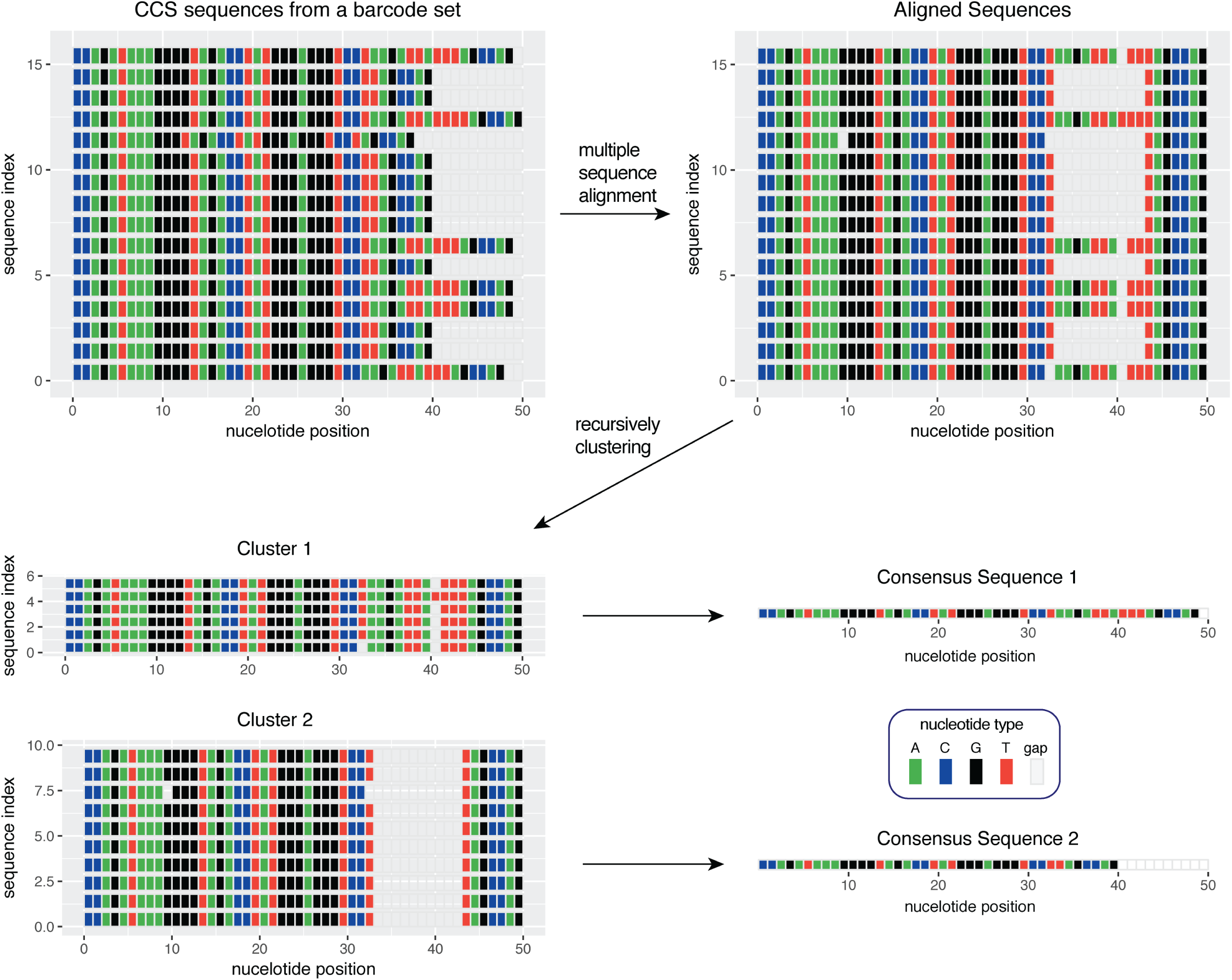
A schematic outline of the PacBio raw data processing. Details are described in Methods.

### PacBio raw reads analysis

To obtain DNA sequences amplified from genomic DNA in single cells, we developed an in-house analysis pipeline (Fig. 2a; the full version is available at https://github.com/Hongyang449/scVDJ_seq). To calculate CCS reads of inserts, PacBio ConsensusTools software version 2.0.0 was used with the following settings: -- inFullPasses=2, --minPredictedAccuracy=90, --minLength=10 and default for other parameters (https://github.com/PacificBiosciences/SMRT-Analysis/wiki/Documentation).

First, to demultiplex CCS reads, we grouped the CCS reads based on the presence of a barcode primer set flanking each end of the CCS. We obtained approximately 35,000 CCSs per PacBio RS II SMRTcell on average, and 46% of them started with a barcode-AF primer and ended with a barcode-AR3 primer. The remaining 54% had mutations, deletions, or additions within the barcoded primer sequence or bore truncated barcode primer sequences, and were excluded from the analysis.

Next, we generated sub-groups in each single cell (unique barcode set) based on the presence of a V or D primer sequence, and sequences in each sub-group were aligned using multiple sequence alignment software MUSCLE(15) (http://www.drive5.com/muscle/). For each group, hierarchical clustering based on sequence length was performed to create initial clusters, then k-means clustering was recursively performed on the DNA sequences of each initial cluster to generate the final multiple clusters, until every nucleotide position in each cluster achieved >51% identity. Clusters containing only 1 CCS read were excluded. Aberrant CCSs with two PCR products ligated together probably resulted during the smartbell adapter ligation step, and were also excluded from the analysis.

To obtain accurate genomic DNA sequence, we calculated a consensus sequence from multiple CCSs in each cluster. Of note, each CCS file that was generated from single zero-mode waveguide (ZMW) corresponds to one copy of PCR product. If a mutation happened during the first PCR cycle, 50% of the final PCR products inherit the mutation at a specific base position in the final PCR product. Similarly, a mutation during the second cycle is inherited by 25% of the final PCR products, 12.5% after the third cycle and so on. After adapter and J region genomic DNA sequences (except the J region used for rearrangement) were removed, the cluster consensus with V primer sequences were submitted to IMGT HighV-Quest analysis(16) (http://www.imgt.org/) to determine whether a specific VDJ sequence in a cluster was predicted to encode a productive or unproductive TCR*β* protein. For example, when DNA sequences rearranged with J1-3, J1-4 to J1-7 regions as well as 3’ of J1-7 to J1 primer sequences were removed. The removal of the extra J region genomic DNA is essential to accurately determine the productivity of rearrangement, especially when some bases are truncated from J regions.

Based on sequence and functionality predicted by IMGT analysis, each cluster consensus sequence was categorized into a tag shown in Supplementary Fig. S1a. Since the status of an allele (GL, DJ, uVDJ and pVDJ) can be determined from possible combinations of 1-2 cluster tags (a total of 24 different patterns are shown in Supplementary Fig. S1b), the rearrangement status from within a single cell (the combination of two separate allelic patterns) was calculated from 2-4 tags recovered from each cell (for a total 253 different combinations of tags). From those, the summary tables were generated.

In this study, the rearrangement status of both alleles was obtained from 909 single cells (25% of 3,664 wells) for thymocytes post beta-selection and 193 DN3a stage thymocytes (25% of 784 wells) when V-to-DJ rearrangement takes place. The frequency depends on sequence depth and was 40% when we recovered 29,643 CCSs from 576 wells. We obtained 0-4 clusters per single cell (1.62 clusters per cell on average) in this depth of sequencing. PacBio CCS data, the barcode list for each sample, genomic DNA and CDR3-IMGT amino acid sequences for all clusters generated from this study are deposited at (https://umich.box.com/s/3f5q64u2i68dn2i7hlneucxph9oetpvb).

## RESULTS

### High-throughput *Trb* genomic DNA sequencing from single cells

To sequence both alleles of genomic DNA in the *Trb* gene VDJ region, we employed a multiplex strategy (Fig. 1a; details described in Methods). Since mouse thymocytes were directly sorted into lysis buffer in individual wells of 96-well PCR plates.

Multiplex, nested PCR reactions followed by barcoding PCR amplified the germline (GL: unrearranged configuration) genomic DNA, D-to-J rearranged (DJ) DNA, and all potential V-to-DJ rearranged (VDJ) DNA in the *Trb* locus(13) (Figs. 1b and 1c). The PCR products were mixed and then sequenced in a PacBio RS II long-read sequencer (www.pacb.com). Each circular consensus specification (CCS) read of an insert generated by the PacBio system corresponds to a single molecule of the original PCR product (Fig. 2a). Based on barcoding and sequence similarity, multiple CCSs were grouped into clusters (Fig. 2a). For each cluster, the consensus sequence was retrieved, which corresponds to an original PCR product. The consensus sequences were subsequently submitted to IMGT HighV-Quest analysis(16) (http://www.imgt.org/) to determine whether the VDJ sequences in any cluster predicted that a functional TCR*β* protein could be generated in that cell. From one *Trb* allele, two [(V-)D1-J1 and (V31-)D2-J2 rearrangement] or one [(V-)D-J2] PCR products are amplified (Fig. 1b and Supplementary Fig. S1). Theoretically, between two and four PCR products (clusters) can be recovered from two alleles in each cell. Details for the computational processing of the PacBio output data to calculate allele rearrangement status are described in Materials and Methods.

To calculate the PCR error rate under our experimental conditions, we took advantage of the unrearranged (germline configuration) genomic DNA corresponding to parts of *Trbd1* to *Trbj1-7* and *Trbd2* to *Trbj2-7* (Fig. 1b) in single bone marrow (BM) cells. From single BM cells we obtained two PCR bands (Fig. 1c), which correspond to the lengths of germline DNA amplified using D*β*1 forward and Jβ1 reverse nested primers (2,672 bp) as well as Dβ2 forward and Jβ2 reverse nested primers (1,896 bp). The accuracy of the resultant sequences was 99.768% for each individual CCS, including both PCR and PacBio sequencing errors. To increase the accuracy, we employed an approach to obtain the consensus sequence from more than 2 CCSs. If we obtain only 2 CCSs per cluster, an error at a position results in 50% identity at the position (ex. An A in 1 CCS and a G in the other CCS); such a cluster was then removed because it is not >51% identical. If we recover 3 CCSs, an error at any position results in a 66.6% identity at the position (e.g. An A in 2 CCSs and a G in 1 CCS). In this case A is the consensus sequence at the position in this cluster. When a consensus was generated from 3 randomly chosen synonymous CCSs, the accuracy increased to 99.991%. These data demonstrate that this method using PCR with Q5® High-Fidelity DNA polymerase (NEB) followed by calculating the consensus sequence from multiple CCSs yields >99.99% accuracy, even after a total of 80 PCR cycles.

### *Trb* VDJ rearrangement status in single thymocytes

To precisely determine the genomic status at the *Trb* loci following VDJ rearrangement in single cells, we analyzed wild type DN4, DP and late DP stage thymocytes, in which *Trb* VDJ rearrangement had already been completed (i.e. post-β- selection). Since DJ joining takes place before the DN3 stage on both alleles, all of the cells analyzed here would be predicted to have undergone at least one productive VDJ recombination event on one allele. We obtained a total of 5,877 clusters and assessed the rearrangement status of both alleles from 909 single cells (Fig. 3). 52% (472) of the single thymocytes were in a [pVDJ (productive VDJ rearranged)/DJ] configuration and 41% (377) were in a [uVDJ (unproductively VDJ rearranged)/pVDJ] configuration, as expected from the 60/40-rule(2). 4% (37) of the post-β-selection thymocytes were in a pVDJ/pVDJ arrangement, which was similar to the 3% dual-TCRβ bearing cells as analyzed using cell surface antibodies(17). Stage-specific data is shown in Supplementary Figs. S2 and S3. The V regions that were most frequently utilized for recombination were V13-2 (Fig. 4 and Supplemental Fig. S4), which is also the most frequently used in inflamed lung epitope specific CD8+ T cells(8). 6,415 V-to-DJ rearranged genomic DNA and complementarity determining region 3 (CDR3) amino acid sequences (4,258 productive and 2,158 unproductive) recovered from single thymocytes in this analysis are available online (see Data Access for detail). Surprisingly, we recovered D2-to-J1 rearranged sequences, although only rarely (Supplementary Fig. S2). At the double negative 3a (DN3a) stage, when V-to-DJ rearrangement takes place, only 9% of the cells had generated a productive (pVDJ) allele [0.5% (pVDJ/GL), 7.2% had generated a (pVDJ/DJ) recombinant and 1.5% were in a (uVDJ/pVDJ) configuration, Supplementary Fig. S3], all of which probably represent cells that were about to develop to the next (DN3b) stage.

**Fig. 3.**
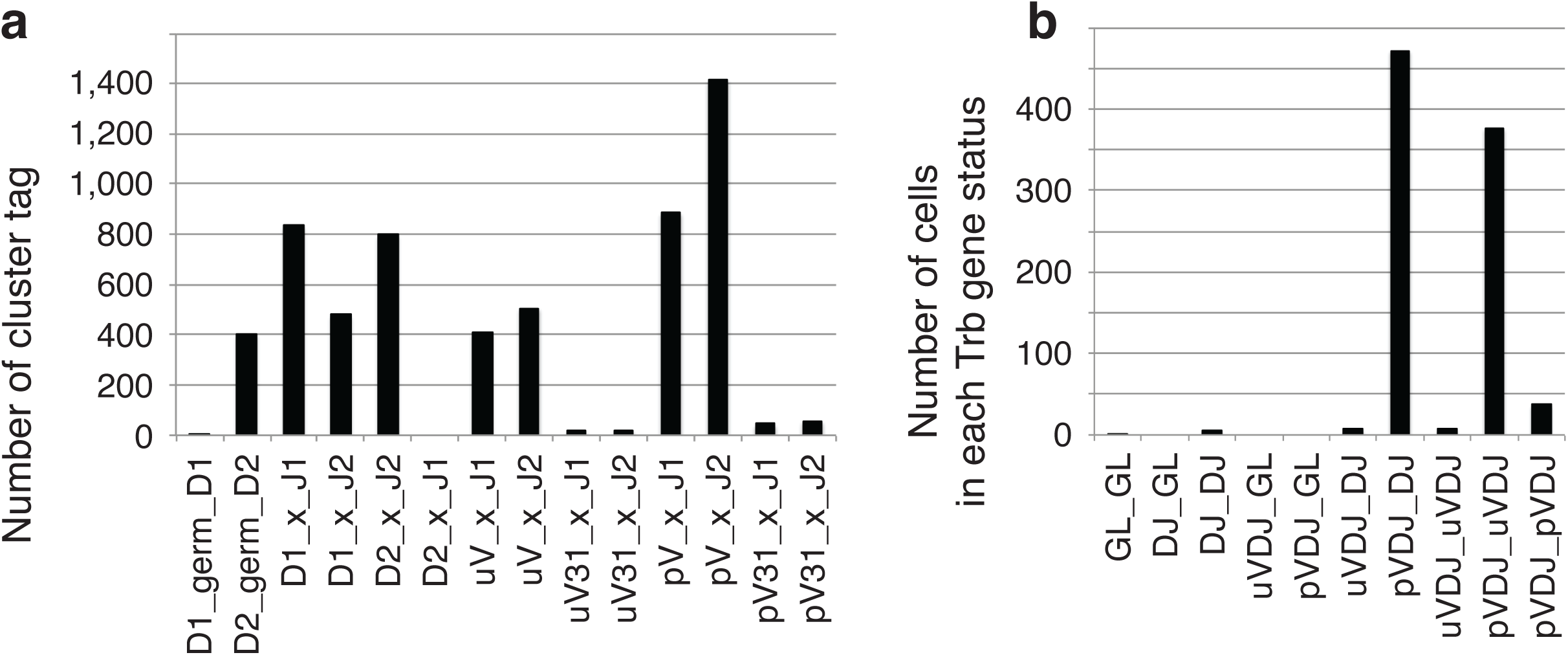
*Trb* VDJ rearrangement status in single thymocytes. (**a**) Cluster tags in a total of 5,877 clusters recovered from single wild type thymocytes at DN4, DP and late DP (post β-selection) stages. Each cluster consensus sequence was categorized into one tag shown on the x-axis (see Supplementary Fig. S1 for detail). (**b**) Calculated *Trb* allele status for 909 single wild type thymocytes at DN4, DP and late DP (post β-selection) stages. Legend: GL: germline allele status; DJ: D-to-J recombined allele; pVDJ: V-to-DJ recombined allele encoding a productive TCRβ protein; uVDJ: V-to-DJ recombined allele encoding a predicted unproductive TCRβ. The data in (**b**) and (**c**) represent a summary of 4 mice for DN4, 5 mice for DP and 4 mice for late DP stages, and details are shown in Supplementary Figs. S2 and S3.

**Fig. 4.**
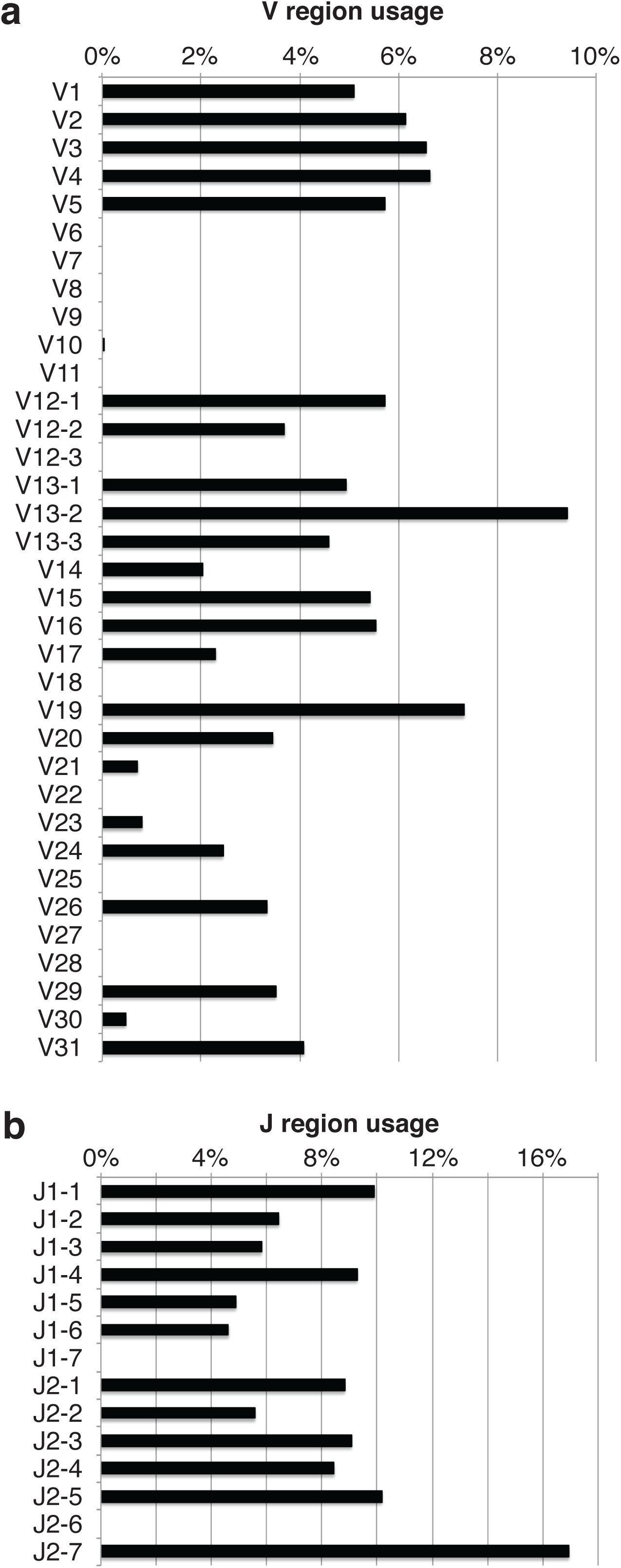
V and J region usage. Frequency of each V region (a) and J region (b) utilization was calculated out of 3,356 VDJ clusters recovered from wild type single thymocytes at DN4, DP and late DP stages. Stage-specific data are shown in Supplementary Fig. S4.

**Fig. 5.**
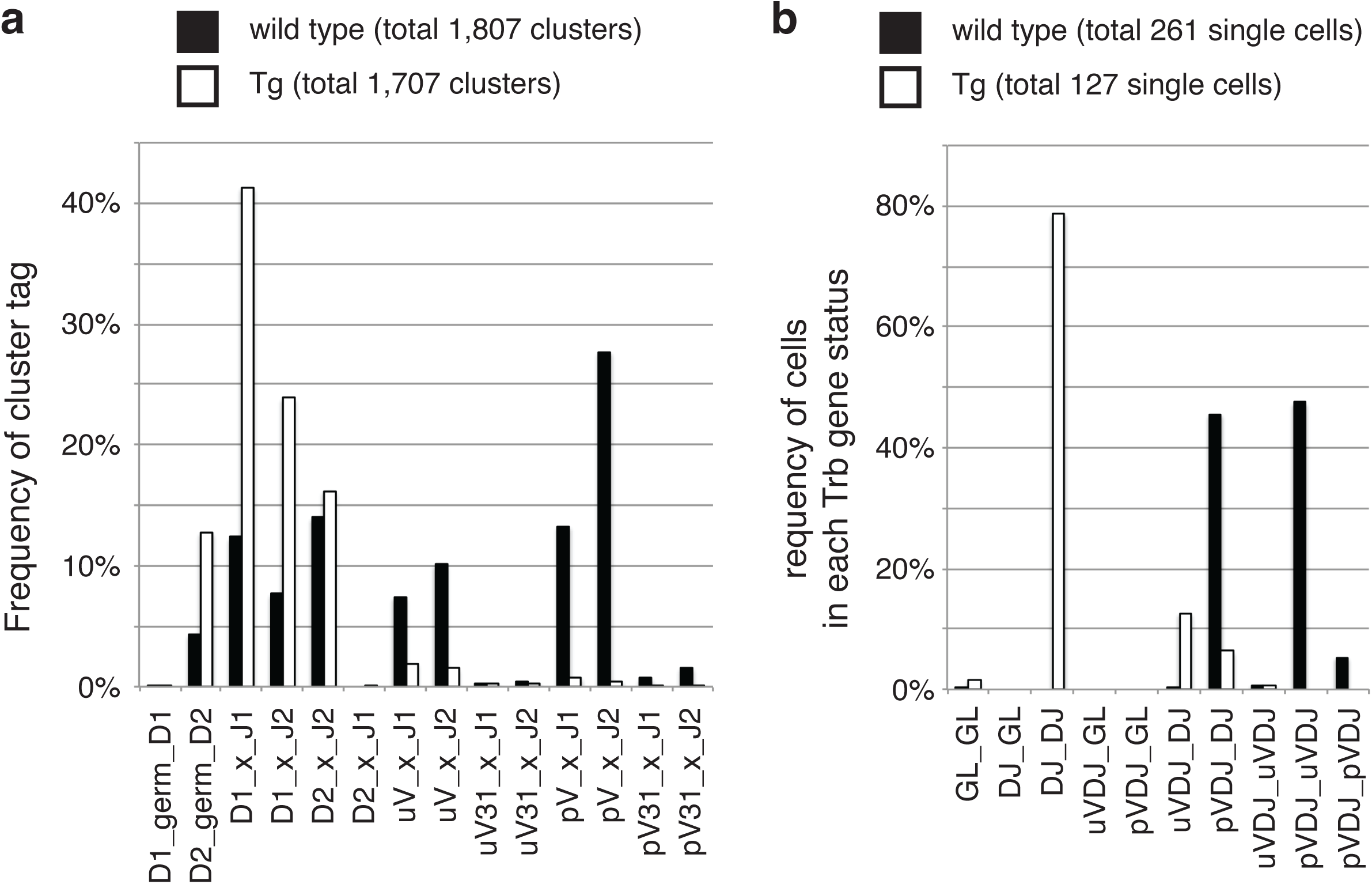
Endogenous *Trb* rearranged status in thymocytes of Vb8 transgenic mice. Number of tags (a) for recovered PCR fragments and rearrangement status (b) calculated from late DP stage single cells isolated from wild type (black) and Vb8 transgenic (Tg, white) mice. Data represent the summary from 4 mice.

Next, to determine whether the methods developed here can capture an artificial allelic exclusion phenomenon, we analyzed late DP cells isolated from Vb8 transgenic mice(18) (Tg^*Vb8*^), which express a productively rearranged TCRβ protein; expression of that transgene has been shown to repress endogenous *Trb* VDJ rearrangement(19). Out of 2,429 clusters generated, 722 clusters bore the DNA sequence of the Tg^*Vb8*^ transgene (Supplementary Figs. S4a) and were removed from further analysis. We then determined the rearrangement status of both alleles from 127 single cells (Supplementary Figs. S4b and S4c). V-to-DJ rearrangement was not observed at the endogenous locus in 80% of the thymocytes, in agreement with published observations(19). However, 20% of the cells had either unproductively (13% uVDJ/DJ) or productively (6% pVDJ/DJ) escaped transgene-suppressed allelic exclusion to allow V-to-DJ rearrangement on one of the endogenous *Trb* loci. In the Tg^*Vb8*^, D2-to-J2 rearrangement was observed less often than D1-to-J1. This supports the idea that D2-to-J2 rearrangement was more frequently repressed when compared to D1-to-J1 region recombination in the presence of a functional TCRβ transgene, in agreement with published analysis of 9 T cell clones(19).

## DISCUSSION

In summary, the method developed here provides an extremely accurate, rapid high-throughput approach for the analysis of *Trb* genomic DNA status at the single cell level. This strategy allowed us to analyze the rearrangement status in thousands of single cells, which is more than 10-fold greater than the number from Sanger sequencing methods previously employed by us and others(10-12) and requiring far less time. This strategy would also be directly applicable to the analysis of the *Igl* and *Igh* genes (encoding immunoglobulins in B cells), which are also regulated by recombination and allelic exclusion at the genetic level(2).

## FOOTNOTES

This work was supported by National Institute of Health Grant AI094642 (to T.H. and J.D.E). The research was also supported in part by the University of Michigan Comprehensive Cancer Center Support Grant (P30 CA046592) for use of the Flow Cytometry and the Sequencing Cores at the University of Michigan.

## AUTHOR CONTRIBUTIONS

T.H. designed the study, performed experiments, wrote computer script, analyzed the data, and wrote the manuscript. H.L. designed the study, wrote computer script and edited the paper. C.K. designed the study, performed experiments, analyzed the data and edited the manuscript. Q.W performed experiments and edited the paper. Y.G. designed the study and edited the paper. J.D.E. designed the study and edited the paper.

## FIGURE LEGENDS

**Supplementary Figure S1.**
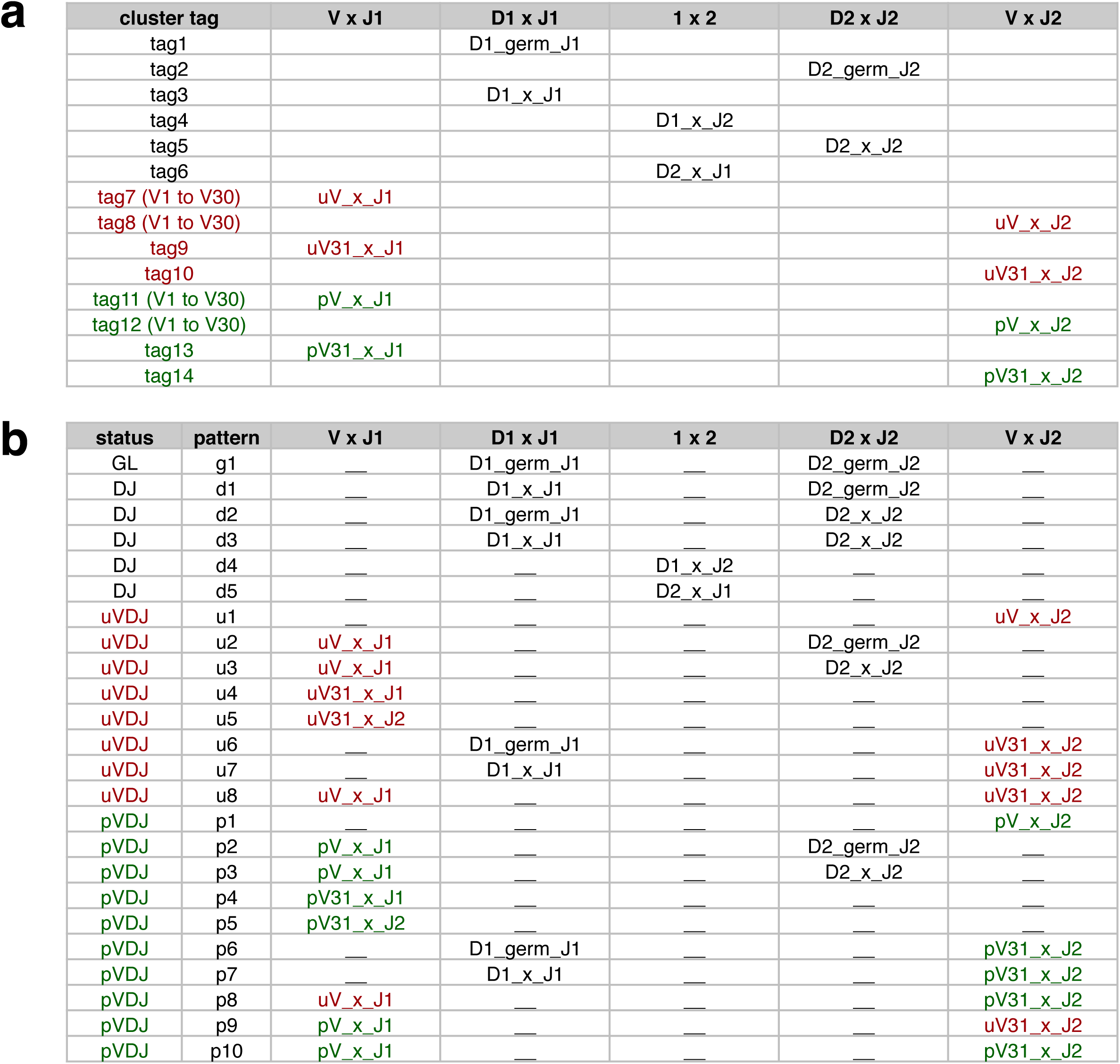
(**a**) Full list of cluster tags (PCR products). PCR fragments amplified using Dβ forward (D1 or D2) and Jβ reverse (J1 or J2) nested primers include both germline configuration (germ) and rearranged DNA (x). PCR fragments amplified using Vβ forward (V) and Jβ reverse (J1 or J2) nested primers encode either productive (p, green) or unproductive (u, red) TCRβ protein. (**b**) Full list of *Trb* gene status and patterns. Status of an allele are shown with possible combination of 1-2 PCR fragments. GL: germline configuration (patter g). DJ: D-to-J rearranged (pattern d). uVDJ: V-to-DJ rearranged DNA encoding unproductive TCRβ (pattern u). pVDJ: V-to-DJ rearranged DNA encoding productive TCRβ (pattern p). Rearrangement with *Trbv31* region (V31) increases the complexity. Full list of possible combination of PCR fragment tags per cell is found in tag_status.csv file in code.zip file. Combination of 2-4 tags used to identify allele status of each cell.

**Supplementary Figure S2.**
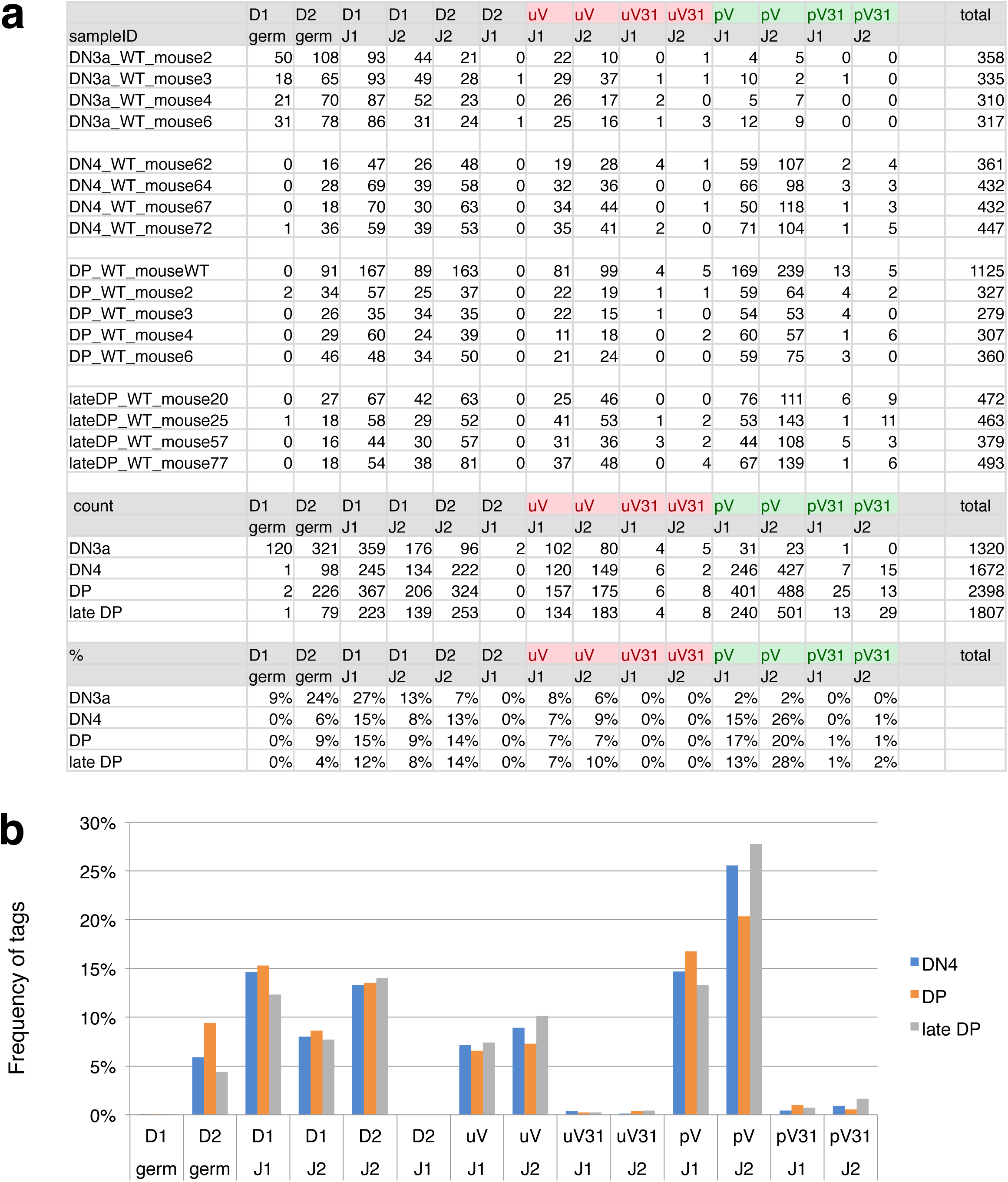
Tags for PCR fragments. Number (**a**) and frequency (**b**) of cluster tags recovered from wild type single thymocytes at DN3a, DN4, DP and late DP stages.

**Supplementary Figure S3.**
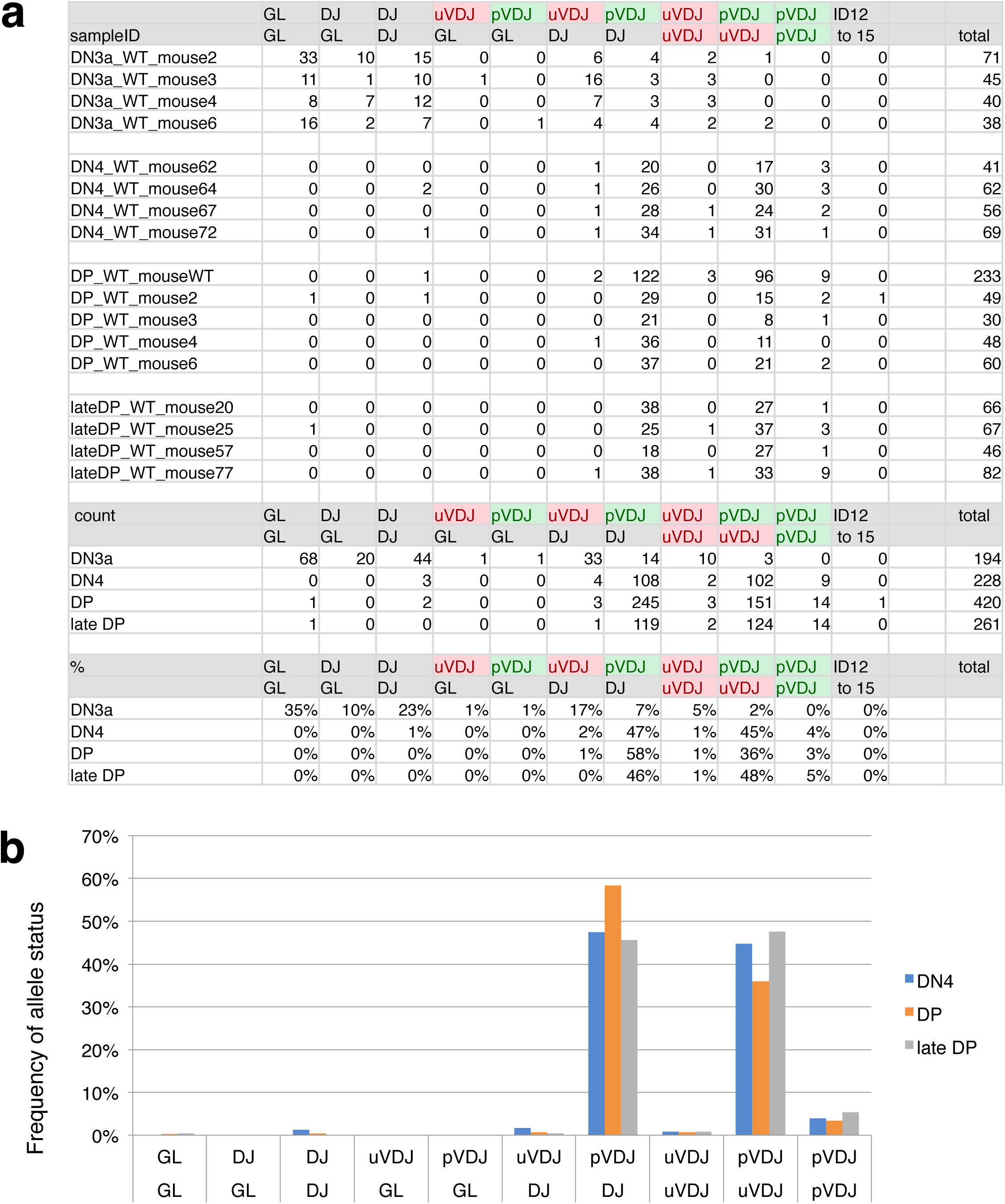
VDJ rearrangement status of both *Trb* alleles. *Trb* VDJ rearrangement status for both alleles calculated from wild type single thymocytes at DN3a, DN4, DP and late DP stages. Number (**a**) and frequency (**b**) are shown. V31

**Supplementary Figure S4.**
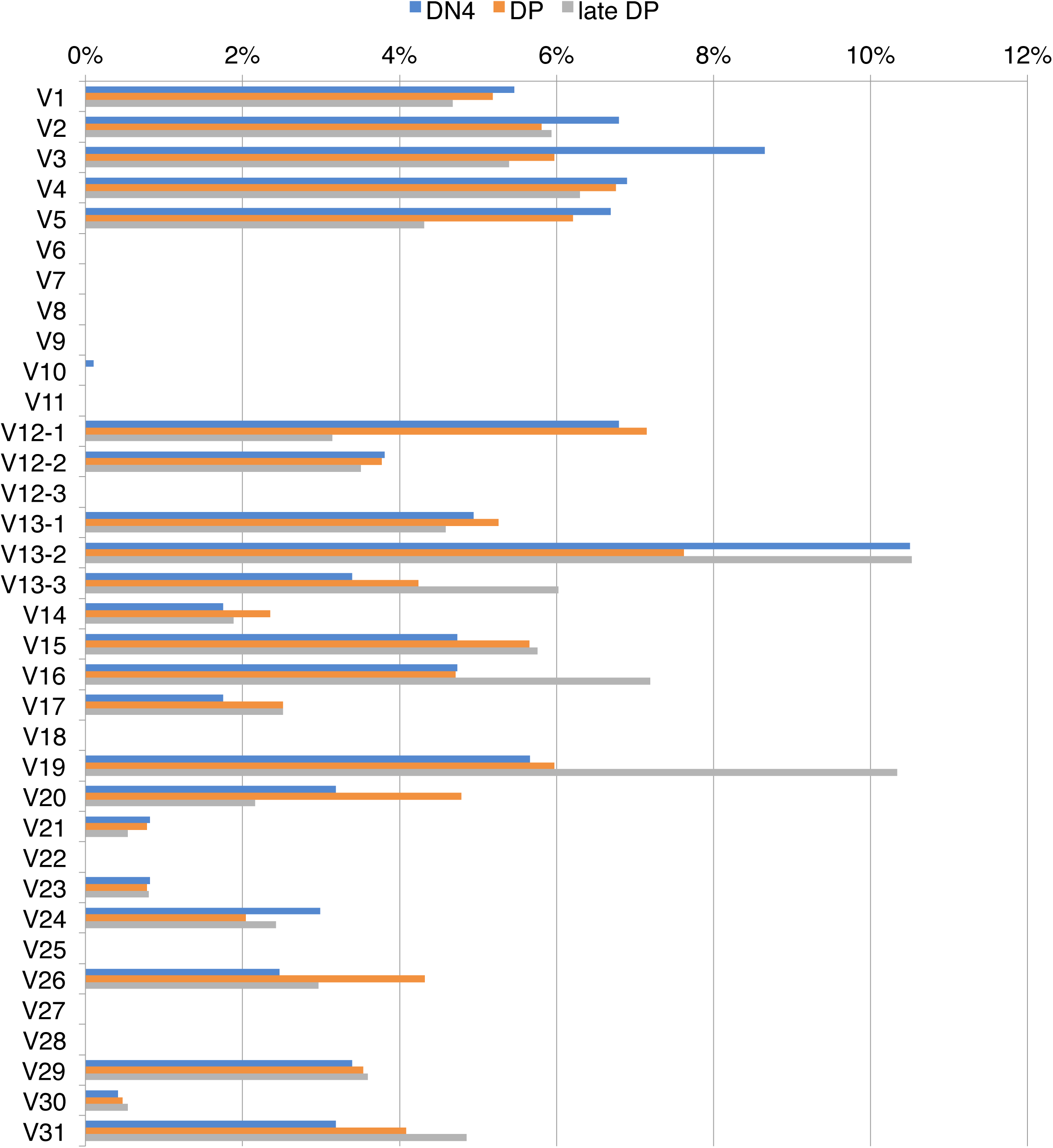
V gene segment usage in wild type single thymocytes at DN4, DP and late DP stages. Frequency was calculated from 971 (DN4), 1,273 (DP) and 1,112 (late DP4) VDJ sequence recovered for each stage.

